# Microplastic pollution in aquatic environments may facilitate misfeeding by fish

**DOI:** 10.1101/2022.07.05.498898

**Authors:** Mitsuharu Yagi, Yurika Ono, Toshiya Kawaguchi

## Abstract

Numerous recent studies have documented ingestion of microplastics (MPs) by many aquatic animals, yet an explanation for misfeeding by fish remains unexplained. Here we tested the hypothesis that biofilm (biofouling) on MP surfaces due to exposure in the aquatic environment facilitates misfeeding in fish. Spherical polystyrene (PS) was cultured for 0 to 22 weeks in a freshwater environment to grow biofilm on the MP. Goldfish were employed in a simple feeding experiment with and without provision of genuine food at ecologically relevant MP concentrations. The absorbance (ABS), which is a proxy for biofilm formation increased exponentially within three weeks of initiation and reached a plateau after approximately five weeks. Although fish did not swallow the MPs, “capture” occurred when food pellets were in the vicinity and significantly increased in probability with exposure time. Duration of capture also increased significantly with increasing exposure. These results suggest that the drift of MPs in aquatic environments may facilitate fish misidentification of MPs as edible prey.

## 1. Introduction

Microplastic (MP) pollution is increasing in all environments globally, including lakes (Dusaucy et al., 2021), rivers (D’Avignon et al., 2021), oceans (Ryan 2015), sediments (Thompson et al., 2004), soils (Zubris and Richards, 2005), and the atmosphere (Panko et al., 2013; Zhang et al., 2020). Since the 1950s, production of plastics has increased exponentially. As of 2015, approximately 5,000 million metric tons (Mt) of plastic, nearly 80% of all plastics ever produced, have accumulated in the environment (Geyer et al., 2017). Approximately 250 Mt of plastics will have accumulated in aquatic environments by 2025 (Jambeck et al., 2015; Wright and Kelly, 2017). These plastics are degraded into small particles due to mechanical stress, photodegradation, and oxidation, instead of decomposing (Eerkes-Medrano and Thompson, 2018). Synthetic polymers < 5 mm in diameter are generally defined as MPs (Andrady 2015; Crawford and Quinn 2016). Most MPs in aquatic environments consist of secondary plastics (Kobayashi et al., 2021) which are derived from the breakdown of larger plastic items via environmental exposure to UV radiation, hydrolysis, wave and wind abrasion, and microbial degradation (Eerkes-Medrano and Thompson, 2018), whereas primary MPs are intentionally manufactured MPs particles, e.g. microbeads in cosmetic products (Alimi et al., 2018; Fendall and Sewell, 2009). This pollution is particularly threatening in aquatic environments, which accumulate MPs long-term due to their very slow degradation (Chamas et al., 2020).

There are global concerns about adverse effects on living organisms via ingestion of MPs. Biota in the aquatic world including bivalves (Baldwin et al., 2020) and other invertebrates (Courtene-Jones et al., 2019), birds (Holland et al., 2016), turtles (Duncan et al., 2019), and fishes (Yagi et al., 2022) ingest MPs. Wotton et al. (2021) reported that approximately 500 fish species ingest MPs, and case reports regarding ingestion of MPs have been increasing. Why do aquatic organisms ingest plastics? Filter feeders such as bivalves and forage fishes obviously are at high risk of feeding on MPs when they are present in the environment (Ding et al., 2021; Tanaka and Takada, 2016). However, mechanisms for selective ingestion of MPs by carnivorous and omnivorous fish are not clear and may be influenced by a multiplicity of factors (Roch et al., 2020). Ory et al. (2017) reported that a planktivorous fish ingested blue MPs as readily as zooplankton. Yagi et al. (2022) also showed that habitat depth and type affect ingestion frequency of MPs in several commercially important fish species. Mechanisms underlying MP ingestion need to be clarified to determine their impact on fish.

As a size matter, MPs have large specific surface area. When pristine MPs are exposed to aquatic environments, their physical and chemical properties change. Specifically, microbial colonisation (biofilms) occurs on MP surfaces, accompanied by deposition of chemical substances such as heavy metals, organic pollutants, and pharmaceuticals (Menéndez-Pedriza and Jaumot, 2020; Pham et al., 2021). These adsorption and adhesion processes have synergistic effects (Johansen et al., 2019; Richard et al., 2019). Biofilms comprise microbial communities including bacteria, algae, protozoa, and fungi, which form multi-layered substrates (Flemming and Wingender, 2010; He et al., 2022; Rummel et al., 2017). The biofilm eventually induces larval settlement of sessile organisms (Bao et al., 2006). In addition, Savoca et al. (2017) reported that forage fish respond to plastic debris solutions, including biofilm. We hypothesize that when MPs are exposed to the aquatic environment, biofilms form on their surfaces, increasing the probability of ingestion by fish. To test this hypothesis, we addressed the following three questions using goldfish, *Carassius auratus*. With increasing exposure time, (i) does the amount of biofilm on the surfaces of MPs increase? (ii) Are fish more likely to ingest them? (iii) Do they accumulate in fish digestive tracts? This study provides insights into MP dynamics in the aquatic environment and their impact upon fish.

## 2. Materials and Methods

### 2.1. Preparation of exposed MPs

MPs for the feeding experiment were made of white, spherical polystyrene (PS), and were originally manufactured as cushioning material (Minibeads, Furukawa Corporation, Tokyo, Japan). In order to reduce particle size variability, stainless sieves of 4 mm and 2 mm were used. MPs exposed to an “aquatic habitat” were prepared by incubating them in the laboratory to prevent their unexpected release into the environment, based on the method of Semcesen and Wells (2021). Water used for culture was from a pond at Nagasaki University, Japan (32º47’08’’N, 129º51’56’’E). The pond is a rectangular tank (10.9 × 6.9 × 0.5 m) made of concrete with well water flowing through it (1.6 L/min). Culturing was accomplished in seven glass containers (900 mL), each containing approximately 4 g of MPs, aerated at 70 mL/min to promote MP submergence. Statistical independence of time series data was ensured by preparing the seven culture containers. Containers were placed near a window in the Fish and Ships Laboratory at Nagasaki University under natural conditions, and MPs were cultured from August 2020 to January 2021. During the incubation period, water in the glass containers was half replaced daily with water from the pond. Pristine MPs were used as Week-0. Temperature Data Loggers (model UTBI-001, Onset Computer Cooperation, MA, U.S.A) were submerged in containers, and water temperature was maintained at 25 ºC with a room air conditioner.

### 2.2. Biofilm quantification

Biofilm biomass on MP surfaces was quantified based on slight modifications of the methods of Lobella and Cunliffe (2011) and Richard et al. (2019). Briefly, five groups of MPs, each with different exposure periods (0, 1, 2, 8, 12 and 18 weeks) were randomly sub-sampled and transferred to 10-mL glass tubes with 1.0 mL crystal violet (0.7% aqueous C_25_H_30_ClN_3_, Hayashi Pure Chemical, Osaka, Japan). After 5 mins at 25 ºC, MPs were washed by vortexing with deionized water until the solution was clear. Then they were dried on filter paper for 10 mins. MPs were transferred to new glass tubes containing 3.0 mL of 95% ethanol and inverted to mix. After 15 mins, 1.0 mL of the solution was measured for absorbance (ABS) at 590 nm using a spectrophotometer (ASUV-3100PC, As One Corporation, Osaka, Japan).

### 2.3. Fish

Fish were purchased from an aqua pet store. Fish were transferred to rearing tanks (0.7 × 0.5 × 0.3 m, 3 replicates of each) filled with 87.5 L of pond water with aeration and acclimated for two weeks before the feeding test. Fish were stocked at a density of 10 – 15 individuals per tank. Water in the tank was changed by half with water from the pond every three days. Water temperature was maintained at 25 ºC, with air conditioning. Fish were fed with artificial food pellets (Kingyonoesa Ootsubu, Kyorin Corporation, Hyogo, Japan) once a day.

### 2.4. Feeding experiment

In this study, the behaviour of taking MPs into the oral cavity is defined as “capture”, and the presence of MPs in the gastrointestinal tract is defined as “swallowing”, according to Ory et al. (2018). All fish were fasted for 24 h before experimental trials. Fish were randomly captured from the rearing tank with a hand net and transferred to a rectangular glass tank (0.4 × 0.3 × 0.2 m) filled with 19.0 L of water. Then, fish were acclimated for 1 h. After acclimation, two trials were conducted to investigate whether fish capture and/or swallow MPs (Fig. 1). For Trial 1, one MP was introduced into the experimental tank with fine tweezers for the fish to eat. The MP was placed in an acrylic cylinder (diameter 40 mm, height 30 mm) located just below the water surface, and the cylinder was suspended from the top of the tank with fishing line (Fig. 1). Trial 1 lasted for 5 min. Individuals that showed no change in behaviour in Trial 1 were advanced to Trial 2. Because the presence of real food makes fish feed actively (Roch et al., 2020), we expected to investigate the frequency and capturing time in Trial 2. In trial 2, three food pellets were added to the cylinder with tweezers. Both MPs and food pellets were floating during the experiment. During trials 1 and 2, fish behaviour was observed and recorded with a digital video camera (Fig. 1).

**Fig. 1.**
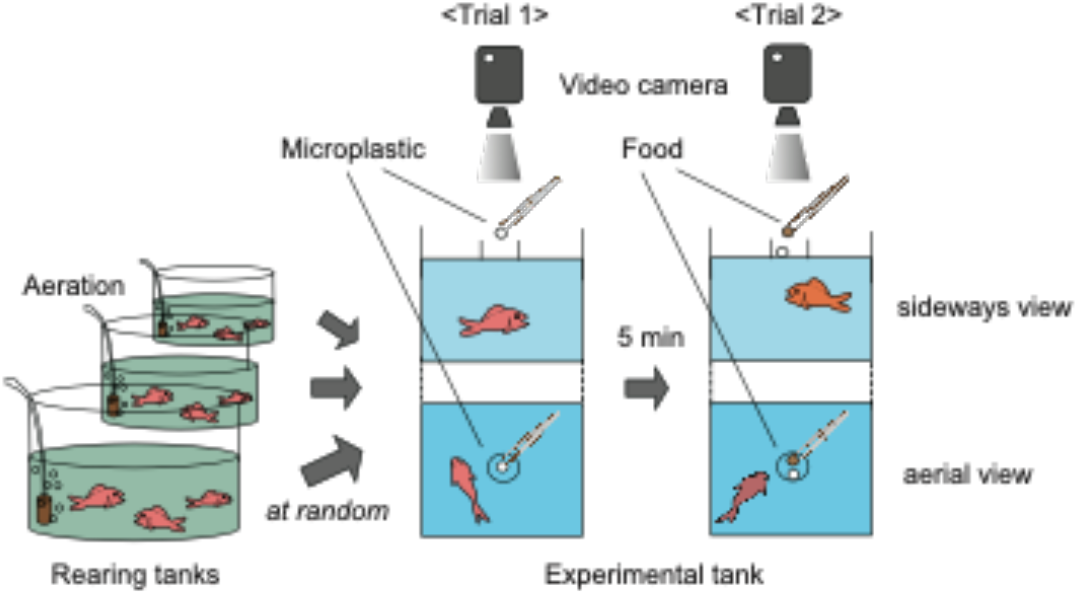
Schematic of experimental design. Fish were chosen randomly from replicate rearing tanks (3) and accommodated in experimental tanks. Fish experienced feeding trials only once, so as to avoid learning-based behavior. In trial 1, fish were provided a single MP particle for 5 min. Subsequently, artificial three food pellets were added (trial 2). Video recordings of fish behavior were made.

The experiment was conducted with one fish per trial and multiple fish (4-12 individuals) using MPs subjected to different exposure periods (0, 1, 13, 14, 17, 21 and 22 weeks). The total number of experimental fish (mean ± s.d.) was 54 individuals (total length: 77.8 ± 8.4 mm, wet weight: 6.6 ± 1.6 g). Fish experienced feeding in the acrylic cylinder for the first time to eliminate effects of learning. Particle diameter means ± s.d. of MPs and food pellets were 3.31 ± 0.40 mm (n = 40), and 2.73 ± 0.19 mm (n = 40), respectively. The water temperature of the experimental tank was maintained at 20ºC with a room air conditioner, and all water was changed after each trial.

All experiments were conducted in accordance with Japanese Animal Care guidelines and approved by the Nagasaki University Fish and Invertebrate Experimental Ethics Committee (Ethics Approval No. NF-0051).

### 2.4. Statistical model

The relationship between exposure period and absorbance (ABS) was examined using a non-linear exponential regression model, i.e., the monomolecular growth model (Fekedulegn et al., 1999), since the surface area of MPs is finite and the amount of biofilm is expected to plateau at a certain time (Richard et al., 2019). For behavioural analysis (trials 1 and 2), we used a generalized linear model (GLM) with a binomial error distribution and logit link function with a binomial response variable of “captured” or “not captured” and exposure period, body size, and interactions as explanatory variables. The relationship between capture time and exposure period was analyzed with the non-parametric Spearman’s rank correlation test, because of their potential nonlinear relationship, including zero value (Suda and Makino, 2016). All statistical analyses were performed using JMP Pro.15, and a probability of p < 0.05 was considered significant.

## 3. Results

Biofilm formed on MP surfaces during exposure experiments, but no macrozoobenthos, such as mussels, were observed. Absorbance (ABS), which is a proxy of biofilm formation, increased exponentially with increasing exposure, within three weeks of initiation, and reached a plateau after approximately five weeks (Fig. 2). The relationship between log (ABS) and exposure period (weeks) could be represented by a non-linear regression (unimolecular growth model) consisting of three parameters (asymptote, scale and increasing rate) (Fig. 2). Estimates ± S.E. were -1.11 ± 0.06 (p < 0.001) for asymptote, -1.40 ± 0.12 for scale, and -0.70 ± 0.13 (p < 0.001) for increasing rate.

**Fig. 2.**
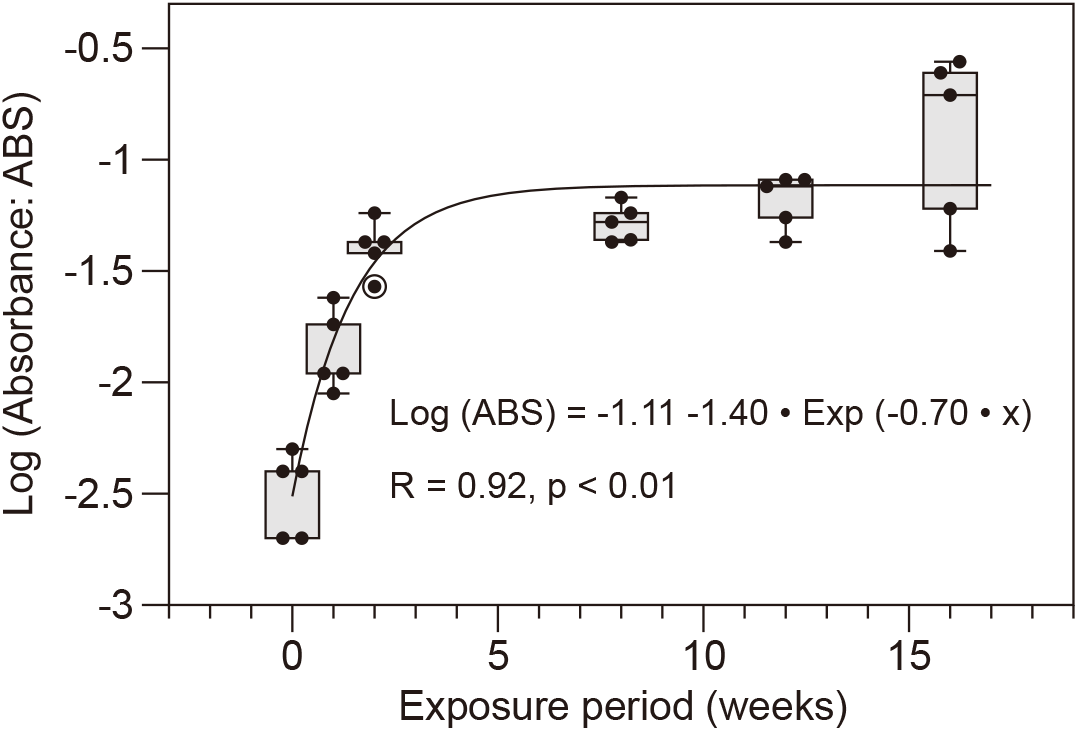
Relationship between absorbance (log ABS) and exposure period (weeks) for MPs. Data are fitted with a nonlinear regression model (molecular growth model). The median (horizontal line), 25^th^ and 75th percentiles (box), and range (bars) of Log (ABS) are presented.

Although “swallowing” did not occur in either trial 1 or 2, “capture” did occur and its probability increased significantly with MP exposure time in Trial 2 (Fig. 3 and Table 1). All individuals captured MPs, spitting them from their mouths, not their gill openings. In Trial 1, a test of the goodness of fit of the whole model compared to a model with only an intercept parameter, was not significant (likelihood χ^2^ = 6.12, df = 3, p = 0.11). Probability of capture in “weeks” was non-significant (p = 0.13), as well as body size (p = 0.79) (Table 1). In Trial 2, the whole model test was significant (likelihood χ^2^ = 12.14, df = 3, p < 0.001) and the “weeks” parameter was significant (p < 0.001), but not body size (p = 0.37) (Table 1). The proportion of individuals that ingested food pellets in Trial 2 was 73% (n = 52), excluding those that captured MPs in Trial 1 (n = 2). The probability of fish ingesting food pellets was not significant in the overall model test (likelihood χ^2^ = 7.65, df = 3, p = 0.05), but the parameter “weeks” was significant (p = 0.02) (Table 1).

**Table 1.** Probability of microplastic “capture” by goldfish increased significantly with exposure time in Trial 2. Estimate, standard error (SE), likelihood ratio Chi-square (χ^2^) and p-value (p) of fixed effects (Exposure time, body size and interactions) on Trials 1 and 2 for MP capture, as estimated by GLM analysis. On Trial 2 for food pellets, parameters are also shown. Significant effects are given in bold.

| Fixed effects | Estimate | SE | $\chi^2$ | p |
| --- | --- | --- | --- | --- |
| Trial 1 |  |  |  |  |
| Intercept | -17.307 | 29.193 | 0.818 | 0.366 |
| Exposure time | 0.428 | 0.517 | 2.277 | 0.131 |
| Body size | 0.851 | 3.307 | 0.070 | 0.791 |
| Exposure time×Body size | -0.041 | 0.347 | 0.014 | 0.905 |
| Trial 2 |  |  |  |  |
| Intercept | -10.351 | 6.248 | 4.396 | **0.036** |
| Exposure time | 0.277 | 0.127 | 11.479 | <<br>**0.001** |
| Body size | 0.625 | 0.684 | 0.798 | 0.372 |
| Exposure time×Body size | -0.049 | 0.073 | 0.434 | 0.510 |
| Food pellet |  |  |  |  |
| Intercept | -1.288 | 2.158 | 0.378 | 0.539 |
| Exposure time | 0.095 | 0.045 | 5.363 | **0.021** |
| Body size | 0.235 | 0.317 | 0.596 | 0.440 |
| Exposure time×Body size | 0.054 | 0.034 | 3.031 | 0.082 |

**Fig. 3.**
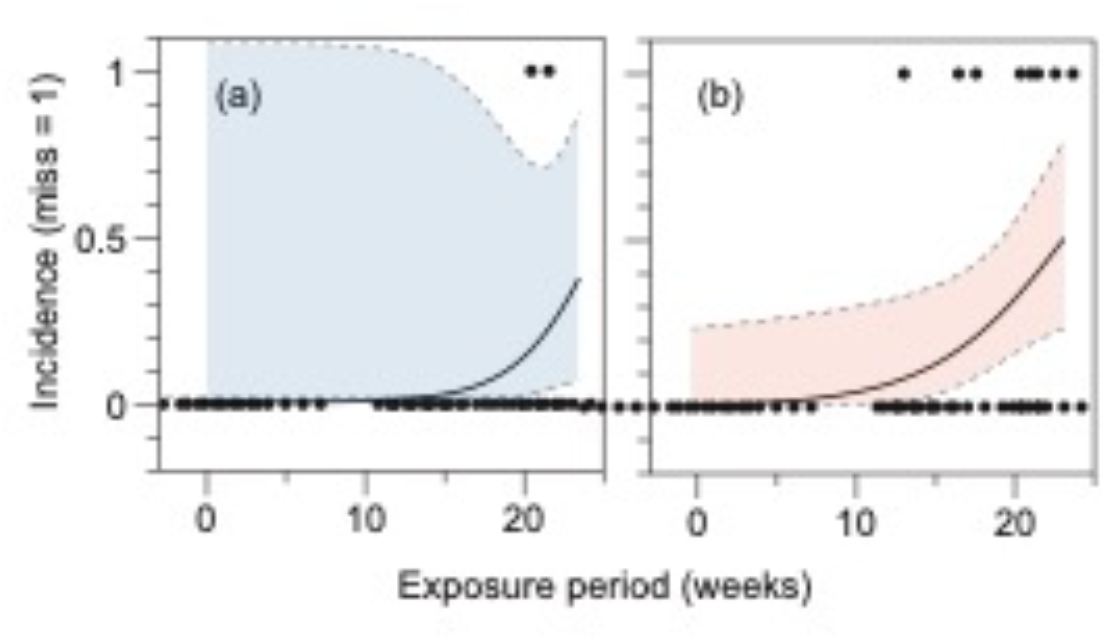
Relationship between the predictive value of incidence (1 = capture occurred) and exposure period (weeks). (a) Trial 1. (b) Trial 2. The 95% confidence interval is shown by the colored area.

Duration of capture increased significantly with increasing exposure period “weeks” (p < 0.001) (Fig. 4). MPs with shorter exposure periods were less likely to be captured, and even if they were, they were spit out in a relatively short period of time. Duration of capture ranged from 24s to 820s.

**Fig. 4.**
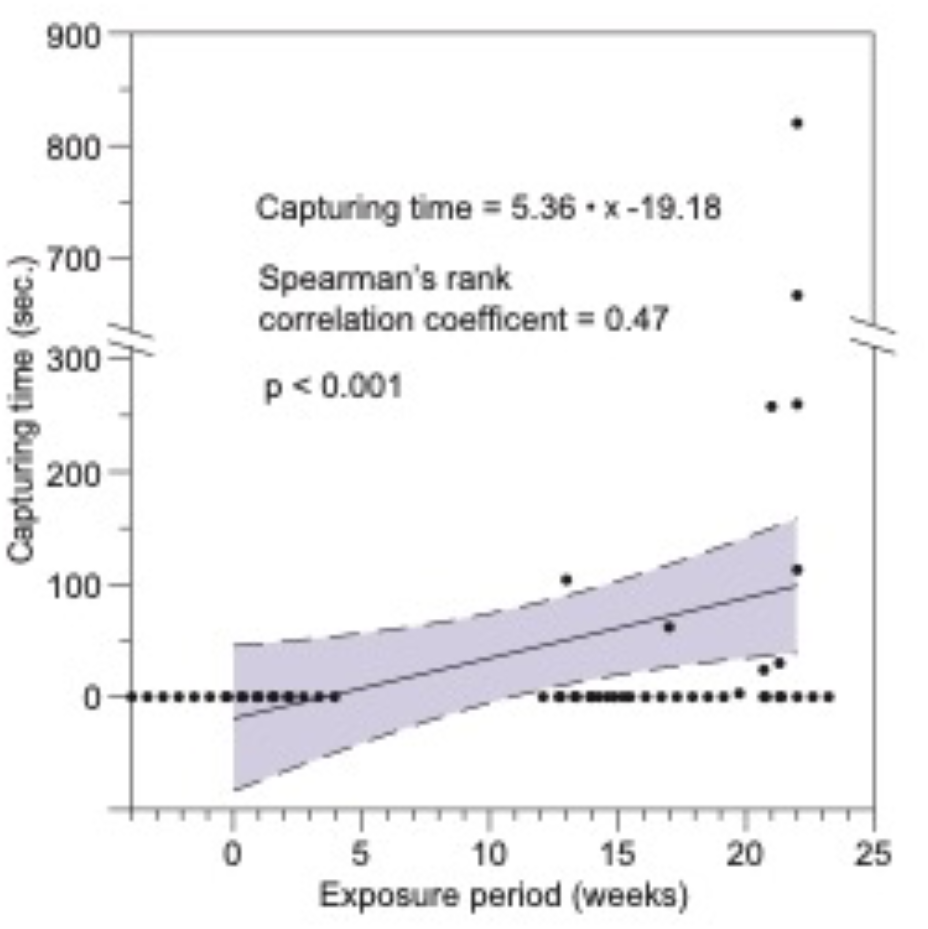
The relationship between capture time (time of MP in the oral cavity) and exposure period. The 95% confidence interval is shown by the coloured area.

## 4. Discussion

To the best of our knowledge, this study provides the first evidence that exposure of MPs to the aquatic environment has a direct effect on fish feeding behaviour. With increasing exposure time, the working hypothesis (i) that the amount of biofilm on the surface of MPs increases, was supported, and (ii) that fish are more likely to take MPs into their mouths, was also supported (Fig. 3). In addition, capture time increased with increasing MP exposure time (Fig. 4). However, the hypothesis (iii) that they accumulate in fish digestive tracts, was not supported, because the fish did not swallow the MPs. Other factors such as particle size and density of MPs, and feeding strategies of fish, was suggested for ingestion of MPs.

Biofilm formation on surfaces of MPs caused physicochemical and biological changes that pristine MPs lack. Pedersen (1982) showed that the amount of crystal violet dye adsorbed to microscope cover slips correlates with the mass of biofilm. Thus, the increase in ABS in this study was due to biofilm formation. In fact, many studies have shown that biofilm formation rapidly occurs in aquatic environments (Parrish and Fahrenfeld, 2019; Rummel et al., 2017), with MPs becoming denser and less hydrophilic (Kaiser et al., 2017; Lobelle and Cunliffe, 2011). Furthermore, biofilm formation triggers accretion of adherent organisms (Zardus et al., 2008). Although we applied a monomolecular growth model, a more detailed observation showed a slight increase in ABS after 10 weeks and a significant increase in ABS in 3/5 of the MPs at 16 weeks (Fig. 2). This is probably due to attachment of other organisms, such as algae, to the biofilm matrix. Indeed, we observed colonization by photosynthetic microalgae, such as diatoms and cyanobacteria. However, we only measured ABS in this study, which limited our assessment of the amounts of adhering organisms. Further studies of effects of chlorophyll and total nitrogen content (Morét-Ferguson, 2010) of MP surfaces on fish misfeeding need to examine longer timescales.

MPs discharged into aquatic environments attract fish. In this study, exposure time did not significantly increase the probability of “capture” in Trial 1, although the p-value was slightly lower (p = 0.13). This is probably due to the smaller attractant effect of MP odor alone. In contrast, a significant increase in exposure time in Trial 2 increased the probability of “capture”, possibly because the presence of food pellets made fish feed actively, and brought the fish in closer contact with the MPs, increasing the probability of intake. This is consistent with the findings of Roch et al. (2020), in which the number of fish ingesting MPs was higher when food was also supplied. Ory et al. (2018) also reported that misfeeding increased when food was present. In addition, odor extracted from biofouled plastic surfaces facilitates food searching behaviour by marine fish (Savoca et al., 2017). It has also been reported that fish tend to ingest MPs of similar color to artificial food pellets (Ory et al., 2017), whereas in the present experiment, capture occurred even though food pellets were brown in colour and MPs were white. The increase in “capture” probability with increasing exposure time may be due to slight differences in color and/or odor, which were not determined in this experiment. Interestingly, food pellet feeding rates decreased significantly with decreasing exposure time (Table 1). We speculate that new, white, shiny MPs may have alerted fish, suppressing pellet feeding.

Increased feeding frequency and time are expected to increase the risk of MP ingestion. A complex of biotic factors, such as fish size, feeding habit, habitat preference, and physical properties of MPs such as particle size, concentration, density, and polymer type may influence ingestion of MPs by fish (Browne et al., 2008; Kumkar et al., 2021; Roch et al., 2020; Rummel et al., 2016; Sathish et al., 2020; Yagi et al., 2022). Goldfish used in this study are visual, omnivores and pick off individual food particles like most diurnal fishes. They actively vacuum up particulate substrate material such as gravel, including food particles. However, they can sort food particles from gravel owing to specialization of the oral cavity (palatal organ) (Finger, 2008). This food-sorting behavior employs taste buds on the palatal organ and the floor of the mouth, and reflex protrusion of the palatal organ to hold food pellets (Finger, 2008). Then, food pellets are moved toward the posterior pharynx (Callan and Sanderson, 2003).

In this study, although fish captured MPs in their oral cavities, they selectively spit out MPs, whereas food pellets were swallowed. Thus, fish detected MPs as foreign particles using taste buds and backwashed MPs to expel them from their mouths. Yagi et al. (2022) reported that in the East China Sea, redwing sea robins, which have sensory pectoral fins, did not ingest MPs, and speculated that fish with taste-detecting organs can avoid MP consumption. Interestingly, capture time in the oral cavity increased with increasing MP exposure duration (Fig. 4). Taste cues from a potential morsel are the primary determining factor for swallowing it (Lamb and Finger, 1995; Sibbing and Uribe, 1985). Thus, the extension of capturing time may be due to the increased time required to recognize MPs as inedible food items, due to biofilm and algae adhesion.

MPs released into aquatic environments have potential adverse effects on fish and fish consumers. Polystyrene (PS) used in this study is less dense than water because it contains air, and it is used in transparent products such as food packaging and laboratory ware (Hwang et al., 2020). PS is a typical microplastic material that is transported over long distances and is commonly found in aquatic environments (Kobayashi et al., 2021). MPs can adversely impact fish health through blockage of and damage to their digestive tracts (Ory et al., 2018), reproductive dysfunction (Sussarelle et al., 2016), reduced food intake (Besseling et al., 2013; Wright et al., 2013), and decreased protein synthesis (Banaee et al., 2020). These deleterious effects may lead to decreased fitness (Fonte et al., 2016; Naidoo and Glassom, 2019). In addition, biofilm on surfaces of MPs lead to sinking and facilitate adhesion of metals, such as Cd, Hg, and Cu (Djaoudi et al., 2022; Richard et al., 2019; Turner and Holmes, 2015), and organic pollutants such as polychlorinated biphenyls (PCBs) and polycyclic aromatic hydrocarbons (PAHs) (Rochman et al., 2013). Ions within biofilms are less tightly bound and easily desorbed (Kurniawan et al., 2012). Biofilm association with MPs may vector these pollutants, enhancing their toxicity (McCormick et al., 2014).

Finally, all MPs used in this study were floating during the experiments, because of their large particle size (∼3 mm in diameter). However, Kaiser et al. (2017) reported that smaller PS particles (∼1 mm in diameter) sink after 14 weeks of incubation, and that the sinking velocity increases by 16% in estuarine water (salinity 9.8). Likewise, Semcesen and Wells (2021) showed that smaller particles have smaller terminal rise and settling velocities than larger particles since the velocities are strongly influenced by drag as well as particle buoyancy (affected by biofilm). In fact, Yagi et al. (2022) reported ingestion of MPs by commercial fishes caught by trawling at 150 m depth. Thus, MPs exposure to aquatic environments appears to increase the risk of MP ingestion Furthermore, prolonged suspension accelerates attachment of larger adherent organisms induced by the biofilm. Although no large organisms were attached in this experiment, we have previously observed large attached organisms on PS plastics that had drifted in the sea for long periods of time (Fig. 5). Attachment of barnacles to plastic fragments has been observed in the North Atlantic Ocean and in bays in South Africa (Fazey and Ryan, 2016; Minchin, 1996). We expect that many fish may selectively feed on plastics to which such large organisms are attached. MP exposure to aquatic environments may facilitate their misidentification by fish as food. Unfortunately, once MPs are released and / or generated in the environment, they cannot be recovered. We urgently need to reduce the amount of plastic that is released into the environment.

**Fig. 5.**
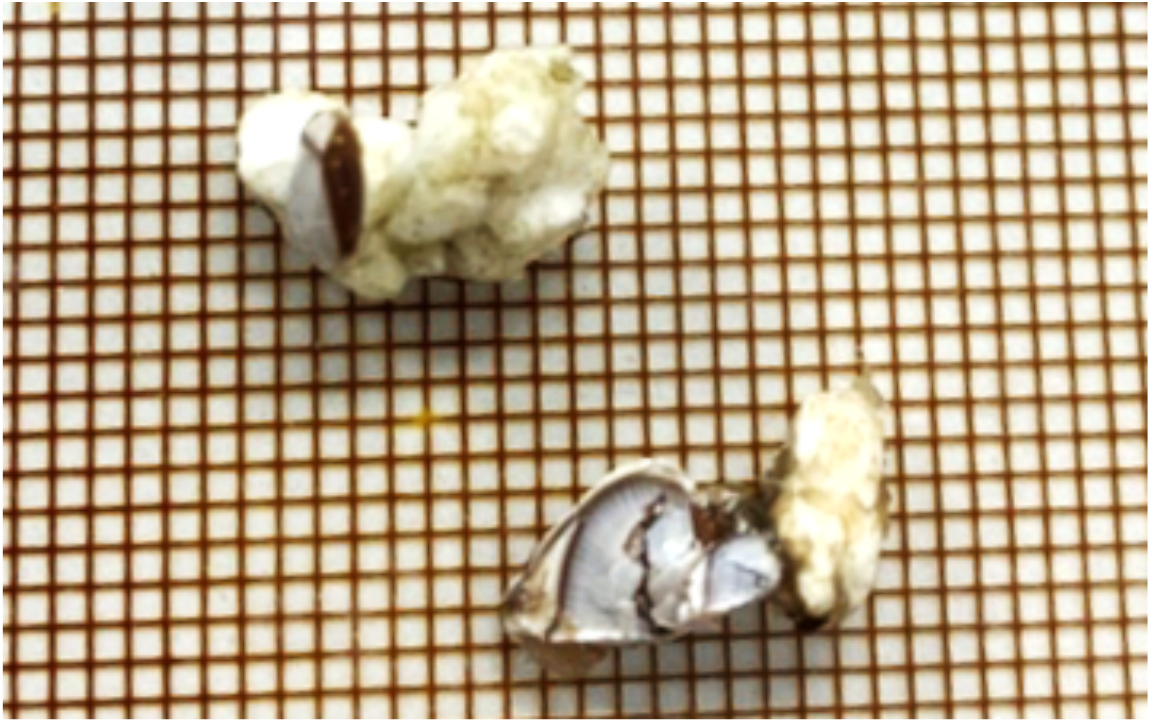
Drifting plastics (polystyrene) and biofouling organisms. Samples were collected in the East China Sea with a neuston net pulled by the T/V Kakuyo Maru, Nagasaki University, in 2020. Mesh size = 1 mm.

## Author contributions

Mitsuharu Yagi: Conceptualization, Methodology, Investigation, Writing – original draft, Supervision,

Funding acquisition, Visualization

Yurika Ono: Investigation, Writing – review & editing.

Toshiya Kawaguchi: Investigation, Writing – original draft, Visualization.

## Acknowledgements

This work was supported by JSPS KAKENHI (Grant Number JP18K14790 and JP21K06337 to M.Y.). We are grateful to students from the “Fish and Ships Laboratory”, Faculty of Fisheries, Nagasaki University, who assisted us with the study. Finally, we thank the editors and anonymous reviewers for their valuable comments and suggestions that greatly improved the quality of the manuscript.

